## Supplementary Information for "Pathway Centric Analysis for single-cell RNA-seq and Spatial Transcriptomics Data with GSDensity"

### Supplementary Notes

#### Supplementary information of the TNBC data analysis.

With the TNBC dataset, we were particularly interested the question whether the tumor cells showed heterogeneity form the angle of their ability of proliferation. We used GSDensity to quantify the heterogeneity of the G2M checkpoint pathway among tumor cells and fetched the actively dividing tumor cells. To generally evaluate the heterogeneity of the G2M checkpoint genes among all 'hallmark' gene sets, we ran GSDensity for all 50 hallmark gene sets. The G2M checkpoint gene set was the 5th most heterogeneous according to the p-values. Moreover, we found that the subpopulation of interest also enriches other hallmark gene sets such as mitotic spindle or glycolysis (Fig 3b-d, p-value < 2.2e-16 for all three gene sets, Chi-squared test), which were all features of actively dividing cells, and thus enhanced our confidence in such observations (not being artifacts). Such enrichments were also observed in TNBC datasets from other patient samples (Extended Data Fig.6d-k).

We also assessed whether such a group of cells could be identified from a routine 'cluster-centric' approach. We found the actively dividing cells appeared as disjoint subpopulations in all four CNV-based clusters (Fig.3f). For transcriptome-based clustering, we clustered the datasets with different 'resolution' parameters to find if there were clusters that enrich the actively dividing tumor cells we identified. We tried to use Silhouette score as a metric to help find a good parameter of the 'resolution' (Fig.3g). With this metric, the best parameter would be 0.1 and result in 2 clusters. For all the resolution tested, under the best case, we found one cluster having only about half (54.5%) of the actively dividing tumor cells (resolution set at 1.2) identified by GSDensity. In conclusion, the actively dividing cells we identified with GSDensity could not be found with cluster-centric approaches. With this example, we demonstrated that GSDensity allowed for knowledge (pathways) guided analysis of limited single-cell data and can effectively generate novel, interpretable and testable hypotheses.

**Supplementary Figures**

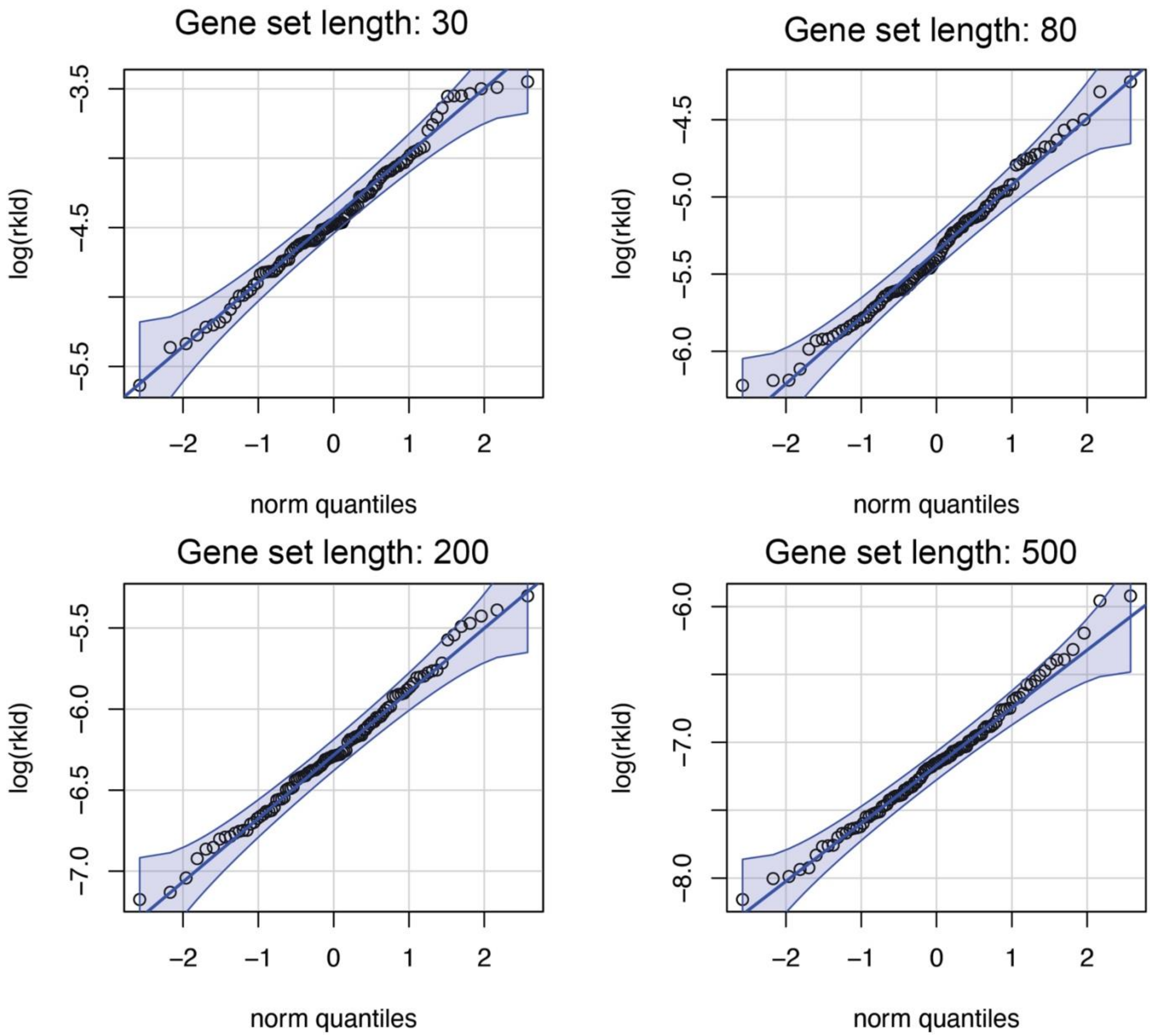


Supplementary Figure 1. The distribution of log-transformed KL-divergence $D_{KL}(P_r||Q)$ between the density of randomly sampled genes and background genes. The four panels showed the qqplot for random gene sets of different lengths, according to the length of most of the publicly curated gene sets.

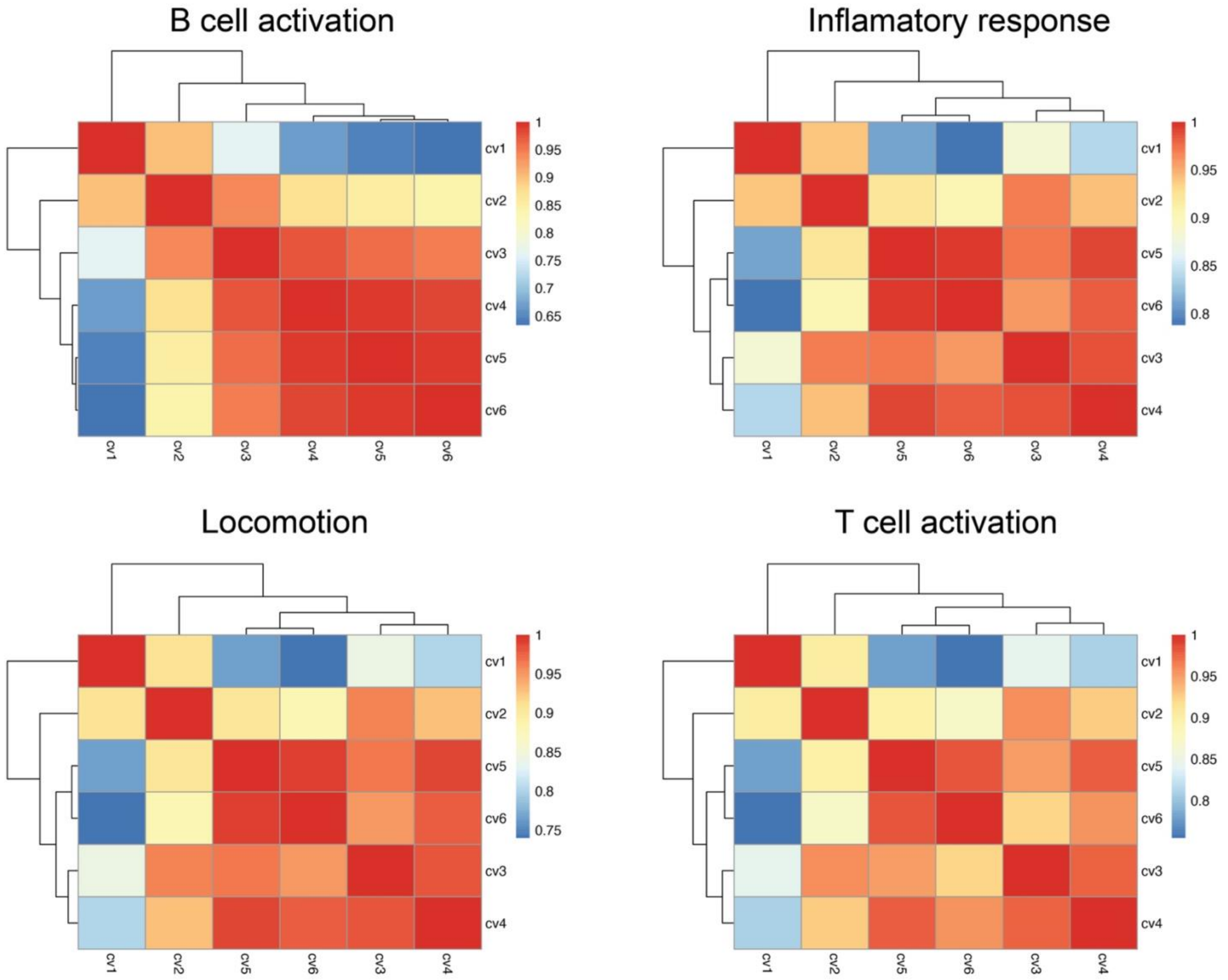


Supplementary Figure 2. The robustness of GSDensity PAL calculation to the parameter (number of neighbors) of the nearest neighbor graph used for network propagation. We show the correlation of single-cell PAL among six different parameters (number of neighbors: cv1: 100; cv2: 200; cv3: 300; cv4: 400; cv5: 500; cv6: 600) for four different pathways in the PBMC dataset.

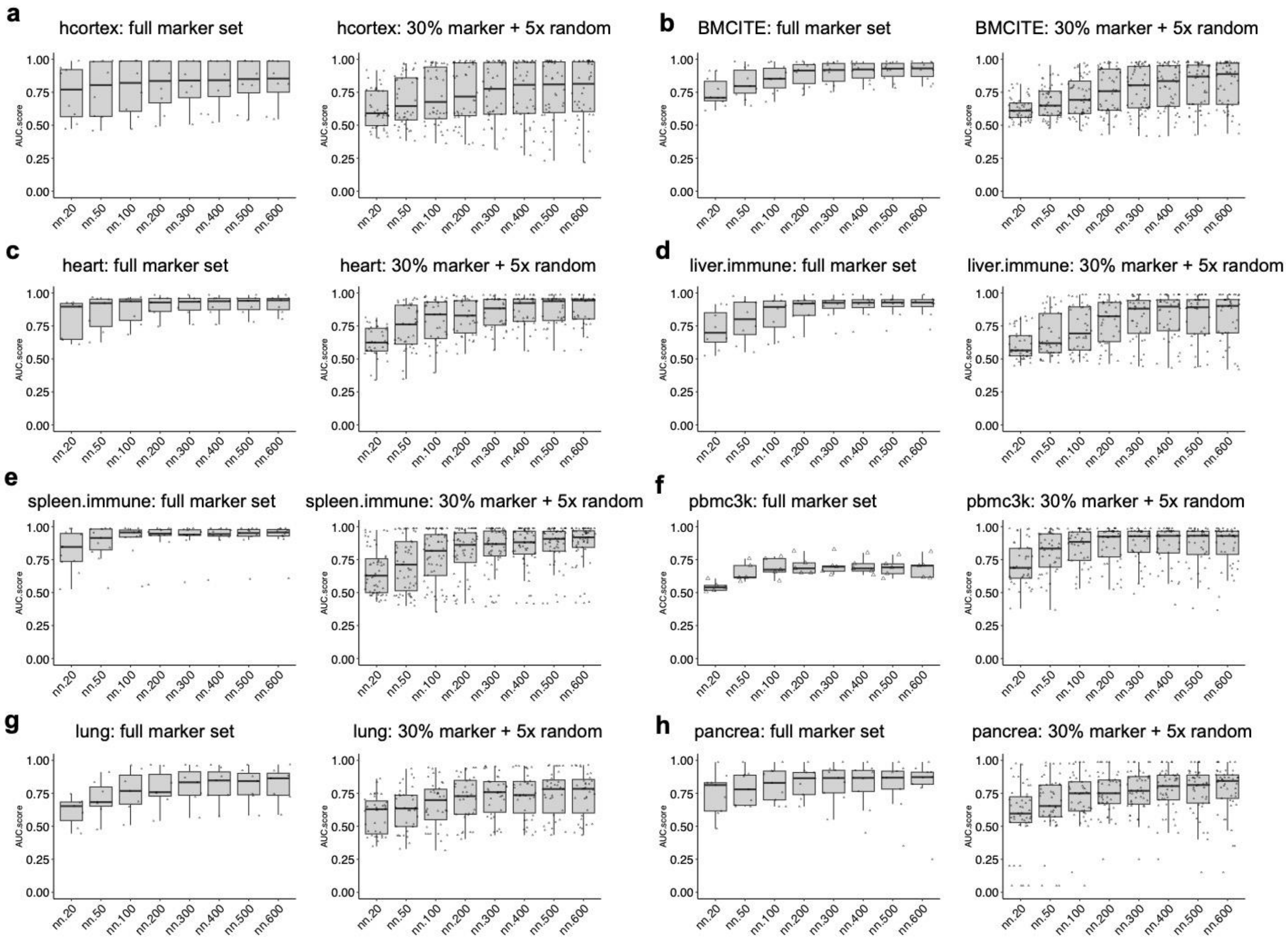


Supplementary Figure 3. The cell identity recovery using marker genes (AUC metric) with different numbers of nearest neighbors used in constructing the NN-graph in GSDensity. For each panel, we demonstrated the recovery with full marker sets (left, strong specificity) or with marker sets mixed with random gene (right, weaker specificity).

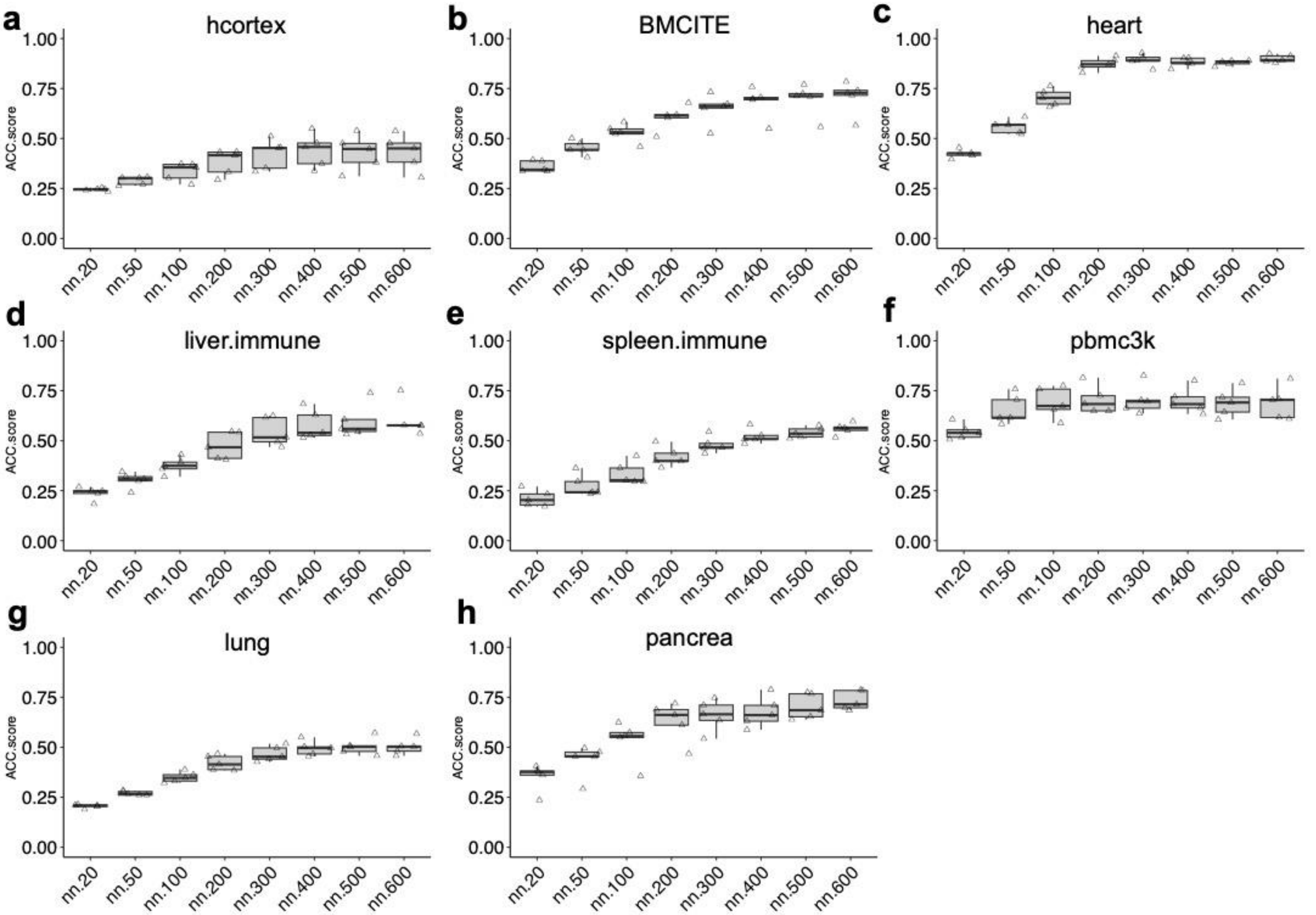


Supplementary Figure 4. The cell identity prediction using marker genes (ACC metric) with different numbers of nearest neighbors used in constructing the NN-graph in GSDensity.

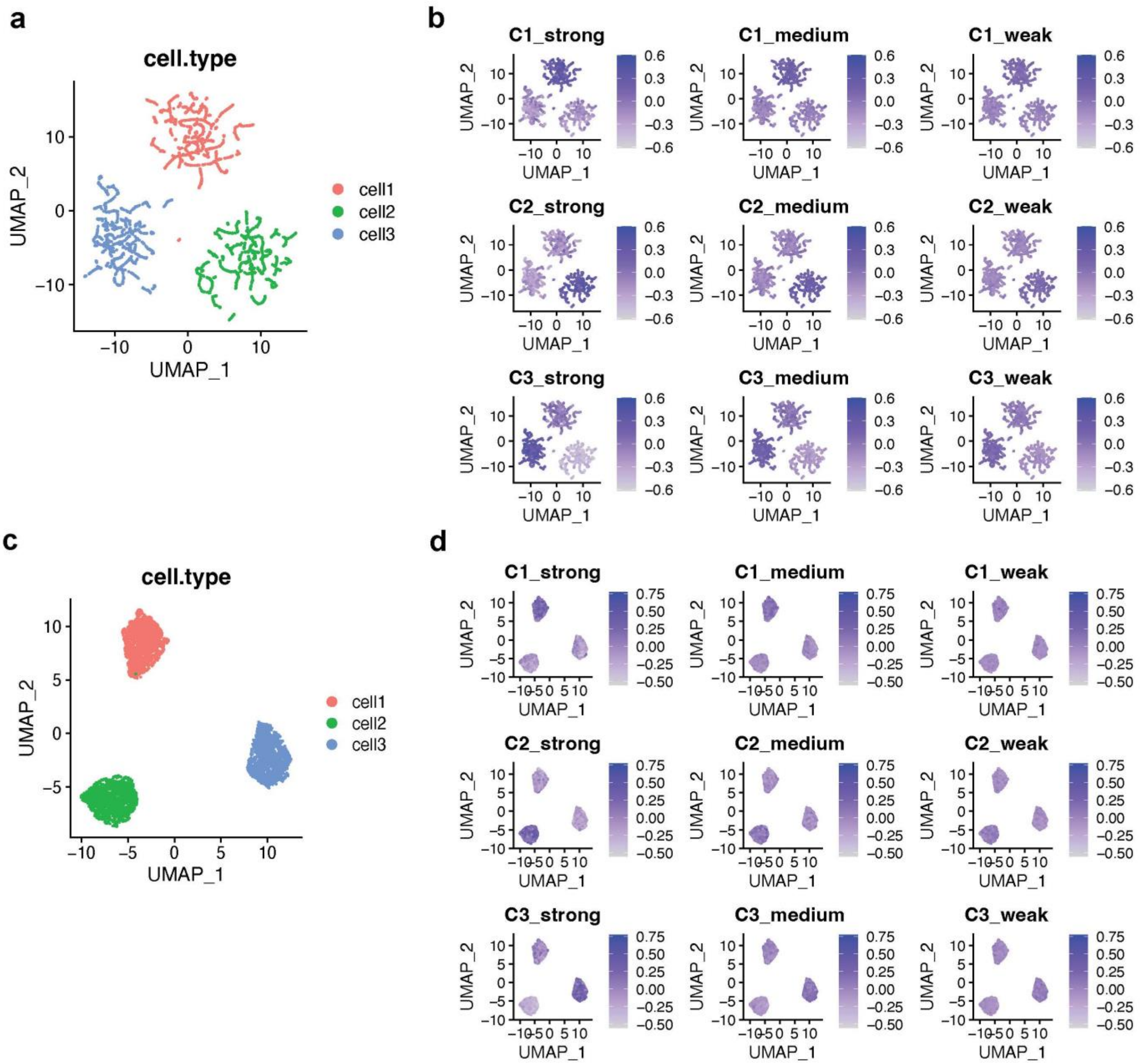


Supplementary Figure 5. Demonstration of simulated scRNA-seq data and marker sets for Mode-1: steady state scRNA-seq data.

a. UMAP visualization of simulated ‘clean’ dataset of Mode-1. Three such datasets were simulated and here we only showed the first batch. The UMAP embeddings were calculated based on gene expression.

b. The overall expression level of strong, medium, and weak marker sets for each cluster for the clean matrix. The expression levels were calculated using the ‘AddModuleScore’ function in Seurat.

c. UMAP visualization of the simulated Mode-1 dataset with noise added. We added noises of different levels to the same clean matrix and resulted in different drop-out rates of the final matrix. Here we only showed the result with 65% drop-outs. The UMAP embeddings were calculated based on gene expression.

d. The overall expression level of strong, medium, and weak marker sets for each cluster for the noise-added matrix. The expression levels were calculated using the ‘AddModuleScore’ function in Seurat.

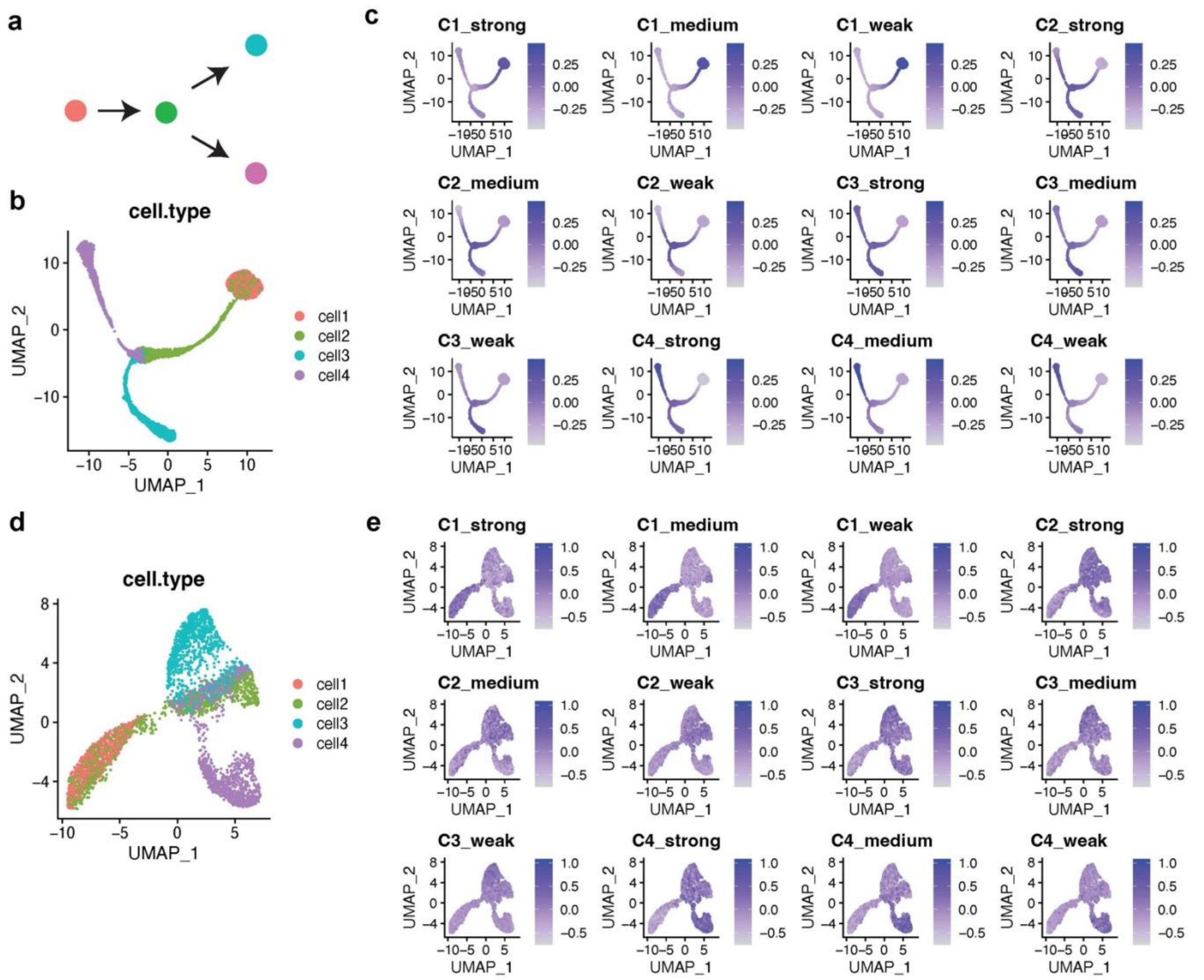


Supplementary Figure 6. Demonstration of simulated scRNA-seq data and marker sets for Mode-2: dynamic scRNA-seq data with four cell states in a bifurcation model.

a. Schematic of the bifurcation model.

b. UMAP visualization of simulated ‘clean’ dataset of Mode-2. Three such datasets were simulated and here we only showed the first batch. The UMAP embeddings were calculated based on gene expression.

c. The overall expression level of strong, medium, and weak marker sets for each cluster for the clean matrix. The expression levels were calculated using the ‘AddModuleScore’ function in Seurat.

d. UMAP visualization of the simulated Mode-2 dataset with noise added. We added noises of different levels to the same clean matrix and resulted in different drop-out rates of the final matrix. Here we only showed the result with 65% drop-outs. The UMAP embeddings were calculated based on gene expression.

e. The overall expression level of strong, medium, and weak marker sets for each cluster for the noise-added matrix. The expression levels were calculated using the ‘AddModuleScore’ function in Seurat.

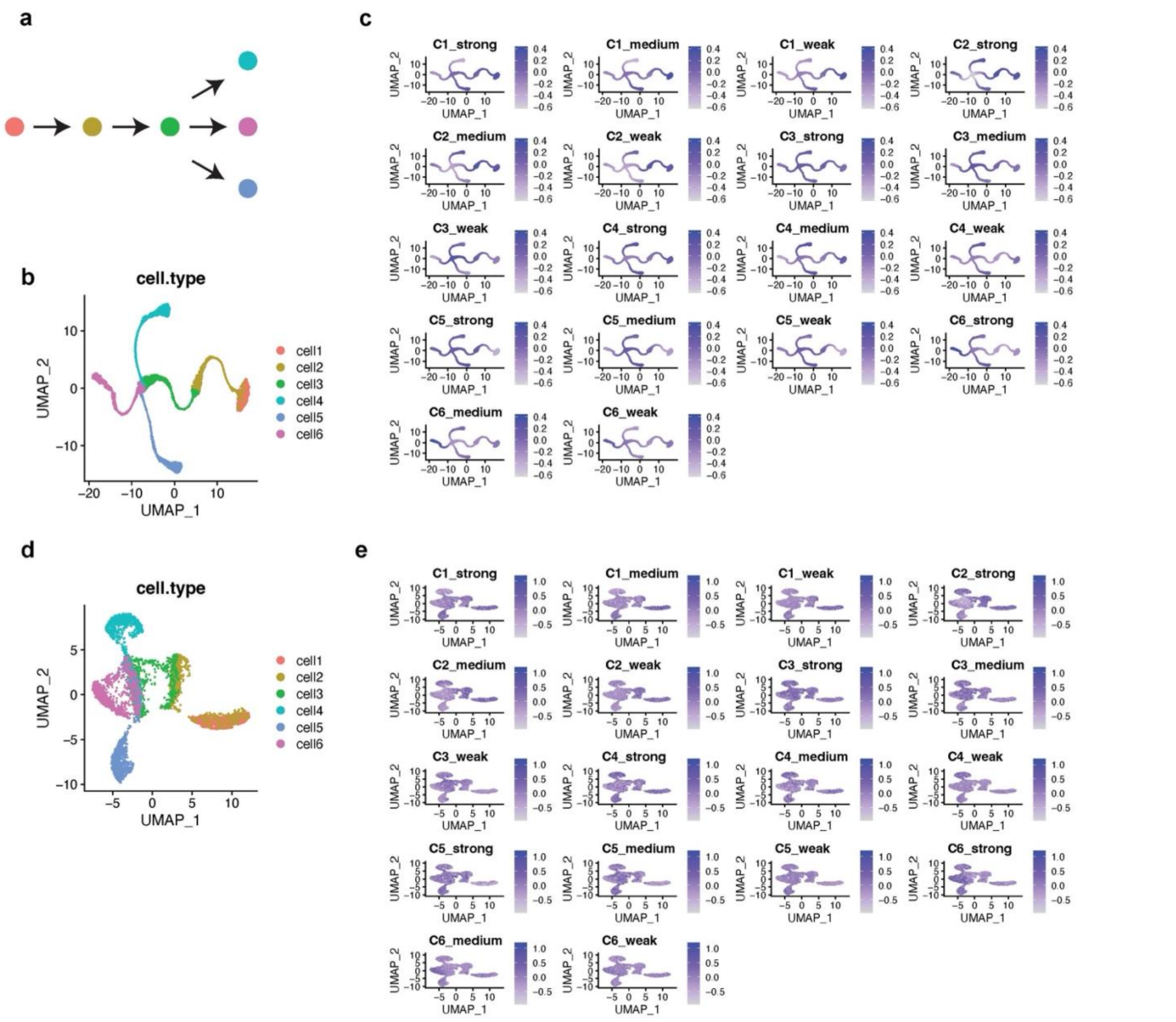


Supplementary Figure 7. Demonstration of simulated scRNA-seq data and marker sets for Mode-3: dynamic scRNA-seq data with six cell states in a trifurcation model.

a. Schematic of the bifurcation model.

b. UMAP visualization of simulated ‘clean’ dataset of Mode-3. Three such datasets were simulated and here we only showed the first batch. The UMAP embeddings were calculated based on gene expression.

c. The overall expression level of strong, medium, and weak marker sets for each cluster for the clean matrix. The expression levels were calculated using the ‘AddModuleScore’ function in Seurat.

d. UMAP visualization of the simulated Mode-3 dataset with noise added. We added noises of different levels to the same clean matrix and resulted in different drop-out rates of the final matrix. Here we only showed the result with 65% drop-outs. The UMAP embeddings were calculated based on gene expression.

e. The overall expression level of strong, medium, and weak marker sets for each cluster for the noise-added matrix. The expression levels were calculated using the ‘AddModuleScore’ function in Seurat.
